# Intraductal Papillary Mucinous Neoplasm Cellular Plasticity is Linked with Repeat Element Dysregulation

**DOI:** 10.1101/2025.01.14.632658

**Authors:** Amaya Pankaj, Michael J. Raabe, M. Lisa Zhang, Yuhui Song, Bidish K. Patel, Katherine Xu, Joshua R. Kocher, Evan Lang, Peter Richieri, Nicholas J. Caldwell, Peiyun Ni, Maria L Ganci, Linda T. Nieman, Vikram Deshpande, Nabeel Bardeesy, Carlos Fernandez-Del Castillo, Martin J. Aryee, Mari Mino-Kenudson, David T. Ting, Yasmin G. Hernandez-Barco

**Affiliations:** Krantz Family Center for Cancer Research, Mass General Brigham Cancer Institute, Harvard Medical School, Charlestown, MA, 02129, USA; Department of Pathology, Massachusetts General Hospital, Harvard Medical School, Boston, MA, 02114, USA; Department of Pathology, Beth Israel Deaconess Medical Center, Boston, MA 02215, USA; Department of Surgery, Massachusetts General Hospital, Harvard Medical School, Boston, MA, 02114, USA; Department of Data Science, Dana-Farber Cancer Institute, Boston, MA, 02115, USA; Broad Institute of Harvard and MIT, Cambridge, MA, 02142, USA; Department of Medicine, Division of Gastroenterology, Massachusetts General Hospital, Harvard Medical School, Boston, MA, 02114, USA

**Keywords:** IPMN, Spatial transcriptomics, cell plasticity, heterogeneity, repeat element

## Abstract

**Background:** Intraductal papillary mucinous neoplasms (IPMNs) are preinvasive pancreatic lesions with a spectrum of histologic phenotypes and variable risk in progressing to invasive cancer. Aberrant repetitive element expression has been shown to be functionally linked to cell state changes in pancreatic cancer.

**Objective:** This study utilized spatial transcriptomics with customized repeat element probes to better understand the relationship of histologic subtypes, repeat element dysregulation, and molecular profiles of different cell populations in IPMN.

**Design:** A total of 52 lesions from 18 patients with resected IPMNs of different histologies and degrees of dysplasia were analyzed with whole transcriptome spatial analysis (GeoMx). Of these, 50 lesions from 17 patients were also processed for single-cell spatial molecular imaging (CosMx). Repeat element probes for LINE1, HSATII, HERVK, and HERVH were used for GeoMx and CosMx.

**Results:** Pancreaticobiliary-type IPMN was enriched for basal-like epithelium and infiltration of Treg cells. Intestinal-type IPMN was enriched for classical epithelium and macrophage infiltrates. Gastric-type IPMN was found to have equal basal-like and classical epithelium with a diverse immune infiltrate. Repeat RNAs were expressed at high levels across IPMN phenotypes and enriched in high-grade dysplasia. Single-cell transcriptional trajectory analysis revealed a phylogeny starting from gastric toward intestinal and pancreaticobiliary branches associated with higher-grade dysplasia and repeat RNA expression.

**Conclusion:** Spatial transcriptomics of IPMN identified a molecular continuum between histological subtypes supporting a common gastric-type origin that transitions to intestinal and pancreatobiliary phenotypes. This cell state plasticity is linked with repeat element expression that can be a potential biomarker for IPMN progression.

**What is already known on this topic:**

- Molecular characterization of IPMN subtypes using regional spatial transcriptomics has described the differences between histologies
- Repeat element expression is associated with cell state changes in pancreatic ductal neoplasm

**What this study adds:**

- Single cell spatial molecular imaging of IPMN subtypes reveals a cellular and molecular continuum starting from gastric histology with distinct branches to Intestinal and pancreatobiliary subtypes
- Repeat element expression is associated with higher grade IPMN histology
- Repeat element expression is elevated in the histologic and molecular continuum from gastric to intestinal and pancreatobiliary-type IPMN

**How this study might affect research, practice, or policy:**

- These results support a common gastric histology origin of IPMN subtypes with different molecular trajectories towards higher grade disease
- Our findings suggest that repeat RNA expression can be used in conjunction with other transcriptional markers of cell state as a biomarker for IPMN progression

## INTRODUCTION

Intraductal papillary mucinous neoplasms (IPMNs) and pancreatic intraepithelial neoplasia (PanINs) are common precursors to deadly pancreatic ductal adenocarcinoma (PDAC) [1, 2]. Whereas PanINs are microscopic and elude detection with standard imaging methods, IPMNs are larger cystic tumors that are detected in 15% of the population, with sharply increased incidence at age >70 years [3]. The likelihood of branch-duct IPMNs developing worrisome features and progressing to invasive cancer varies significantly, ranging from 4.8% to 48.8% depending on size and the presence of worrisome features [3]. Although there have been improvements in clinical and imaging criteria for risk stratification of patients with IPMN, a significant portion of patients still undergo surgical resection that would not have progressed to invasive cancer. Pancreatic surgery has greatly improved, but there is still the risk of major morbidity 14.4% and mortality 0.9% as reported recently from our institution [4]. The combination of increasing IPMN detection and the highly heterogenous outcomes of patients highlights the need to obtain a more comprehensive molecular understanding of these lesions. Conventional histological evaluation of IPMNs encompasses a spectrum of morphologies including gastric, pancreaticobiliary, and intestinal epithelial subtypes. In parallel, histological determination of high-grade dysplasia (HGD) and low-grade dysplasia ( LGD) helps provide prognostic information, but the ability to do so pre-operatively on cyst fluid cytology remains a clinical challenge [5]. Promising diagnostic biomarkers from cyst fluid, pancreatic fluid, and blood are actively being developed, but there remains an interest in identifying novel biomarkers to apply to these biospecimens [5, 6]. Although there have been improvements in clinical and imaging criteria for risk stratification of patients with IPMN, a significant portion of patients still undergo surgical resection that may not have progressed to invasive cancer.

Given the histological heterogeneity of IPMN, there has been interest in determining if there are differences in the microenvironment. There have been extensive studies in PanINs that describe activation of inflammatory pathways, but with defects in anti-tumoral immunity [7, 8, 9, 10, 11, 12, 13, 14, 15, 16, 17, 18, 19, 20]. Notable features include a striking absence of CD8+ cytotoxic lymphocytes (CTLs) and early infiltration of Foxp3+ T-regulatory (T-reg) cells, macrophages, and myeloid-derived suppressor cells (MDSCs) which are considered tumor-promoting immune cells. Studies in IPMN have been more limited, but some studies have described the evolution of the immune microenvironment showing infiltration of T and B cells in low-grade IPMNs with progression to increased Tregs and macrophages in high-grade IPMNs/invasive carcinoma arising in IPMN [21, 22, 23, 24]. In addition, the contribution of stromal fibroblasts and their different phenotypes with primarily myofibroblast and inflammatory cancer-associated fibroblasts (myCAFs and iCAFs, respectively) has been well described in PDAC [25, 26, 27], and some studies in IPMN have shown the presence of CAFs associated with HGD and progression to invasive disease [28, 29].

Here, we have expanded upon these studies with GeoMx analysis of 52 IPMN samples from 17 patients combined with CosMx single-molecule imaging of 1,000 genes in 50 samples from 17 patients from the same cohort. Using both technologies we have characterized transcriptional signatures and trajectories that support the transdifferentiation to intestinal and PB-type IPMN morphologies with HGD from a precursor gastric-type IPMN with LGD. In addition, we have incorporated customized probes designed to detect repeat RNA sequences, which we previously identified as being elevated in human PanIN and IPMN lesions (33, 34).

These repeat RNA sequences have recently been shown to play a functional role in driving critical cell state changes in both cancer cells and cancer-associated fibroblasts (CAFs) within human PDAC (35). We find that these repeat RNAs are dysregulated in high-grade IPMN and track with transdifferentiation from low-grade gastric IPMN, which presents a potential new biomarker of progression from IPMN to invasive carcinoma. By including these repeat RNA probes, we aim to further explore their contribution to tumor progression and the tumor microenvironment in preinvasive lesions.

## METHODS

### Specimen Acquisition and Histological Review

This discarded excess tissue study was approved by the Institutional Review Board at the Massachusetts General Hospital (IRB Protocol 2013P001854). Formalin-fixed paraffin-embedded (FFPE) tissues from patients with pancreatic surgical resections diagnosed as IPMN were used for this project. We identified 52 distinct IPMN lesions from 18 patients, and representative hematoxylin and eosin (H&E) slides were reviewed by gastrointestinal pathologists (MM-K, NJC, MLZ) for tissue microarray (TMA) construction. Three total TMAs were constructed; each consisted of 24-3 mm cores with one core each of normal pancreas and normal lung as control. The presence of lesional tissue in each TMA core was confirmed by pathologists (MM-K, NJC) prior to immunohistochemistry. Another 11 tissue sections from a cohort of four patients who had undergone resection for IPMN were obtained for full slide analysis to evaluate intratumoral heterogeneity. Each whole section had areas of LGD and HGD annotated by gastrointestinal pathologists (VD, BKP). Additional slides were used for spatial transcriptomics and validation as below. Pertinent demographic and clinical data are summarized in Supplementary Tables 1 and 2.

### GeoMx^TM^ Digital Spatial Profiler (DSP)

We used the GeoMx^TM^ Digital Spatial Profiler (DSP) platform (Bruker/Nanostring) to collect genetic information from transcribed mRNA. This involved staining the slide with the GeoMx^TM^ Human Whole Transcriptome Atlas (a mix of unique oligo-labeled probes against 18,676 mRNAs). This was followed by staining with fluorescent morphology markers. The morphology panel included 1:10 SYTO13 (Nanostring/Bruker), 1:20 anti-panCK–Alexa Fluor 532 (Nanostring/Bruker), 1:100 anti-CD45–Alexa Fluor 594 (Nanostring/Bruker), and 1:100 anti-αSMA–Alexa Fluor 647 (clone 1A4, Novus Biologicals, cat. no. IC1420R) diluted in blocking buffer W (NanoString/Bruker). The anti-panCK and anti-CD45 antibodies were supplied pre-conjugated, whereas anti-αSMA was conjugated using the Alexa Fluor 647 Antibody Labeling Kit (abcam, ab269823). Post staining, multiple regions of interest (ROI) were marked out with H&E as a reference. A total of 143 ROIs were marked out on the 3 TMAs, following which the instrument accurately segmented the ROIs into distinct immune/epithelial/fibroblast “areas” of interest (AOI). This was followed by the collection of the barcodes (attached to oligos by ultraviolet light cleavable bonds) onto a 96-well collection plate. The sampled oligos were then pooled following the NanoString GeoMx-NGS Readout Library Prep User Manual (MAN-10117-03). Illumina’s i5 x i7 dual indexing system was used for amplification and ligation of the adapters. Quantitation of final library was done by qPCR using KAPA estimation (KAPA code: KK4854, Roche catalog: 07960298001) using Roche LightCycler® 480 program and eventually sequenced with a 1%PhiX spike-in on a NextSeq 1000/2000 using a 2 x 100 cycle MidOutput v3 flow cell following instructions on the Illumina NextSeq manual (15048776 v16). The generated FASTQ files were processed to generate target probe-specific count files and saved in DCC format.

### CosMx™ Spatial Molecular Imager (SMI)

For this study, we utilized the CosMx™ Spatial Molecular Imager (SMI) from Bruker Technologies to perform a single-cell transcriptomic analysis on three IPMN TMAs. Sample preparation followed the standardized workflow detailed in the CosMx™ SMI Manual Slide Preparation for RNA Assays (MAN-10184-04), which includes deparaffinization, target retrieval, tissue permeabilization, and overnight hybridization with RNA-specific probes. We used the Human Universal Cell Characterization Panel (1000-plex, RNA) with an additional set of custom probes capturing retrotransposons including long interspersed nuclear element-1 (LINE1) retrotransposon open reading frames (ORF1 and ORF2), HSATII, and two human endogenous retroviruses (HERV-K and HERV-H). For cellular segmentation, we employed DAPI Nuclear Stain, CD298/B2M, and PanCK/CD45 as markers. The processed slides were imaged on the CosMx SMI platform, which combines cyclic in situ hybridization and high-resolution fluorescence imaging to detect multiplexed RNA transcripts at single-cell resolution. Field of view (FOV) selection was performed to capture representative regions of interest within the TMAs, including areas with diverse cellular compositions and stromal interactions with varying degrees of dysplasia, among the three histological subtypes. Data analysis was conducted using the AtoMx™ Spatial Informatics Platform (SIP), enabling spatial visualization and analysis of transcriptional heterogeneity and retrotransposon expression within the IPMN microenvironment.

### GeoMx Computational Methods

Probes were collapsed to the geometric mean of each target after removing outliers detected through a Grubbs test conducted on a per-gene basis across areas of Interest (AOIs). AOIs that exhibited low read alignment quality, low total read counts, or insufficient segment area were excluded from the final analysis to ensure data reliability. The data was quantile normalized such that each AOI has the same 75th percentile of expression. For clustering, AOIs were grouped using the Unweighted Pair Group Method with Arithmetic Mean (UPGMA), applied to all genes that displayed a standard deviation greater than 15. This clustering approach helped identify biologically meaningful patterns, and AOIs were subsequently classified by their predominant cellular composition into three key categories: Epithelial, Immune, and Fibroblast segments, which served as the basis for further downstream analyses.

Moffitt metascores and fibroblast metascores were calculated to evaluate the molecular characteristics of different subtypes. Specifically, the Moffitt metascores were determined by calculating the difference in mean z-score between the basal-like and classical subtypes, while fibroblast metascores were computed as the difference in mean z-score between inflammatory cancer-associated fibroblasts (iCAFs) and myofibroblastic cancer-associated fibroblasts (myCAFs).

For differential expression analysis, a Mann-Whitney U-test was applied to compare expression levels, with p-values being FDR-adjusted when appropriate. Gene Set Enrichment Analysis (GSEA) was performed using the GSEA R package version 1.2. Additionally, immune cell subtype composition within the ROIs was assessed using the Spatial Decon R package version 1.6.0, allowing for a detailed characterization of the immune microenvironment.

Scripts and code were written for R and are available on request.

### CosMx Computational Methods

For quality control, 32 out of 390 fields of view (FOVs) were excluded from the analysis due to significant signal loss, primarily caused by high tissue autofluorescence. Individual cells were removed from the dataset if they exhibited low transcript counts (fewer than 20 transcripts per cell) or fell below a predefined size threshold of less than 25 pixels. Cell classification was done using the Insitutype v2.0 algorithm in R. The algorithm was run in an unsupervised approach, using unbiased clustering of cells based on their gene expression profiles. Dimensionality reduction techniques were employed for further analysis. Principal component analysis (PCA) was performed using the irlba R package (version 2.3.5.1). Uniform manifold approximation and projection (UMAP) was then applied using the uwot R package (version 0.2.2). To quantify the activity of gene programs across cell populations, metascores were calculated as the mean z-score of gene expression within each defined program. Additionally, a repeat score was computed for each cell, defined as the sum of the expression levels of repetitive element-associated genes to provide a measure of repeat element activity. Pseudotime trajectory analysis was carried out using the Slingshot R package (version 2.4.0).

Scripts and code were written for R and are available on request.

## RESULTS

### Characterization of the Subtype-Specific IPMN Microenvironment with Whole Transcriptome Digital Spatial Profiling

We utilized the GeoMx platform with the Whole Transcriptome Atlas panel, comprising an oligonucleotide probe set targeting 18,676 genes, to investigate the spatial organization of compartment-specific gene expression programs between the histological subtypes of IPMN. After quality control, our TMA cohort for analysis included 51 formalin-fixed, paraffin-embedded (FFPE) clinical specimens from 17 treatment-naive, resected IPMNs. A multi-step approach was employed to capture specific AOIs corresponding to epithelial, immune, and fibroblast compartments within the IPMN microenvironment. Oligonucleotide probes were hybridized to target mRNAs and fluorescently labeled antibodies against Pan-CK (epithelial), CD45 (immune), and Alpha-SMA (fibroblast) were utilized to segment these compartments. Following segmentation and QC, 101 ROIs were identified across the 51 samples, with a median of 2 ROIs (range 1-3) per core (**Figure 1A**). A surgical pathologist with expertise in pancreatic pathology (MM-K) evaluated each ROI for histological grade and sub-type, enabling compartment-specific profiling of epithelial, immune, and stromal regions (**Figure 1B**). The cohort was distributed across histological subtypes, including 48 gastric, 29 intestinal, and 27 pancreaticobiliary ROIs. Normalized gene expression profiles from each AOI were subjected to unsupervised hierarchical clustering analysis of all genes with standard deviation >15, demonstrating a clear distinction between the epithelial and stromal compartments (**Figure 1C**). Restricting the analysis to epithelial compartments revealed further organization of expression profiles by histological subtype, distinguishing gastric, intestinal, and PB IPMN (**Figure 1D**).

**Figure 1.**
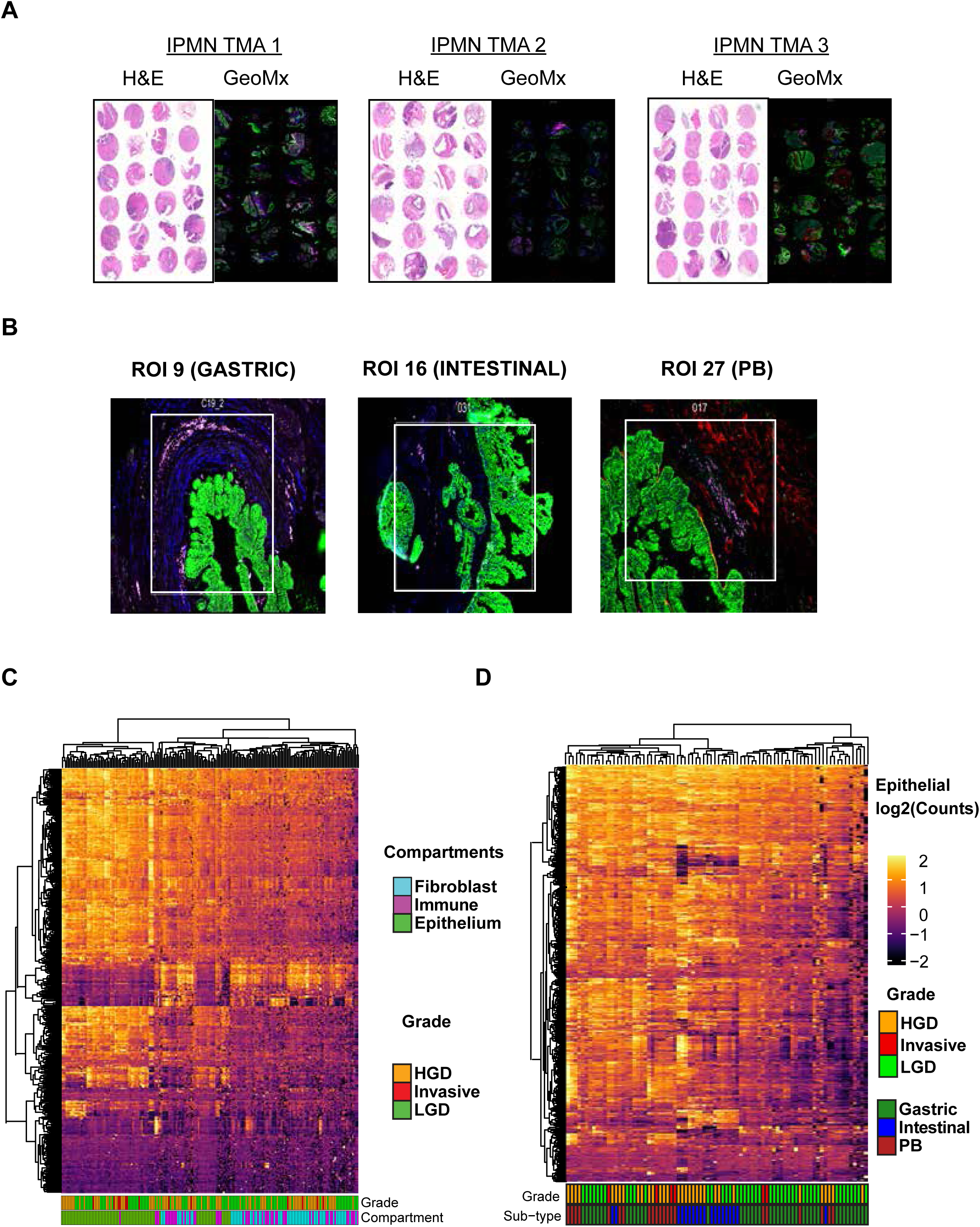
Characterization of the subtype-specific IPMN microenvironment with whole transcriptome digital spatial profiling (A) Representative Hematoxylin and Eosin (H&E) stained cores from the TMA cohort used for digital spatial profiling. Cores were sourced from 17 treatment-naive, resected IPMN patients. (B) Histological examples show three distinct intraductal papillary mucinous neoplasm (IPMN) subtypes. For each example, the H&E image is shown adjacent to the GeoMx visualization of segmentation masks used for Region of Interest (ROI) collection. ROIs were identified corresponding to the epithelial, immune, and fibroblast compartments within the IPMN microenvironment. Representative ROIs for a gastric subtype (ROI 9), an intestinal subtype (ROI 16), and a pancreaticobiliary (PB) subtype (ROI 27) are illustrated. (C) Heatmap visualization of normalized log2(Counts) gene expression profiles (standard deviation > 15) across all collected ROIs (n=101). Unsupervised clustering separates the ROIs into two primary groups corresponding to the epithelial compartment and the stromal compartments (Immune and Fibroblast). (D) Unsupervised hierarchical clustering analysis restricted to the epithelial ROIs (n=48 gastric, n=29 intestinal, and n=27 pancreaticobiliary) reveals clear separation of expression profiles corresponding to the three major histological subtypes of IPMN (Gastric, Intestinal, and PB). IPMN histological grade is shown in the top bar (LGD: Low-Grade Dysplasia, HGD: High-Grade Dysplasia, Invasive).

### The Gastric Subtype – A Heterogenous Transitionary Cluster

Unsupervised hierarchical clustering of all gastric epithelial AOIs (n=42) did not reveal distinct clusters by grade (**Figure 2A**). To identify gastric subtype-specific epithelial cell phenotypes,we used Moffitt gene signatures for classical/epithelial and basal/mesenchymal phenotypes previously defined for PDAC and calculated a meta score defining each epithelial AOI into classical or basal-like for all 42 epithelial AOIs. This analysis identifies that 48% of AOIs are classical, and 52% of them are basal-like with a heterogenous expression of both cell states on the heatmap (**Figure 2B**). To investigate the immune microenvironment of our Gastric cohort, we proceeded with an immune deconvolution analysis to determine the immune cell composition of the sub-type. The gastric IPMN subtype exhibits a diverse immune profile, highlighted by monocyte enrichment (10%), significant CD4 T memory cells (11%) and CD8 T memory (12%) (**Figure 2C**). We used a metascore derived from myCAF and iCAF gene signatures annotated in PDAC to characterize the fibroblast niche. This analysis classified 70% of fibroblast AOIs in the gastric subtype as iCAF and 30% of them as myCAF (**Figure 2D**). In summary, the gastric subtype predominantly exhibits a heterogenous epithelial cell phenotype and microenvironment enriched with diverse, active immune cells and iCAF fibroblasts

**Figure 2.**
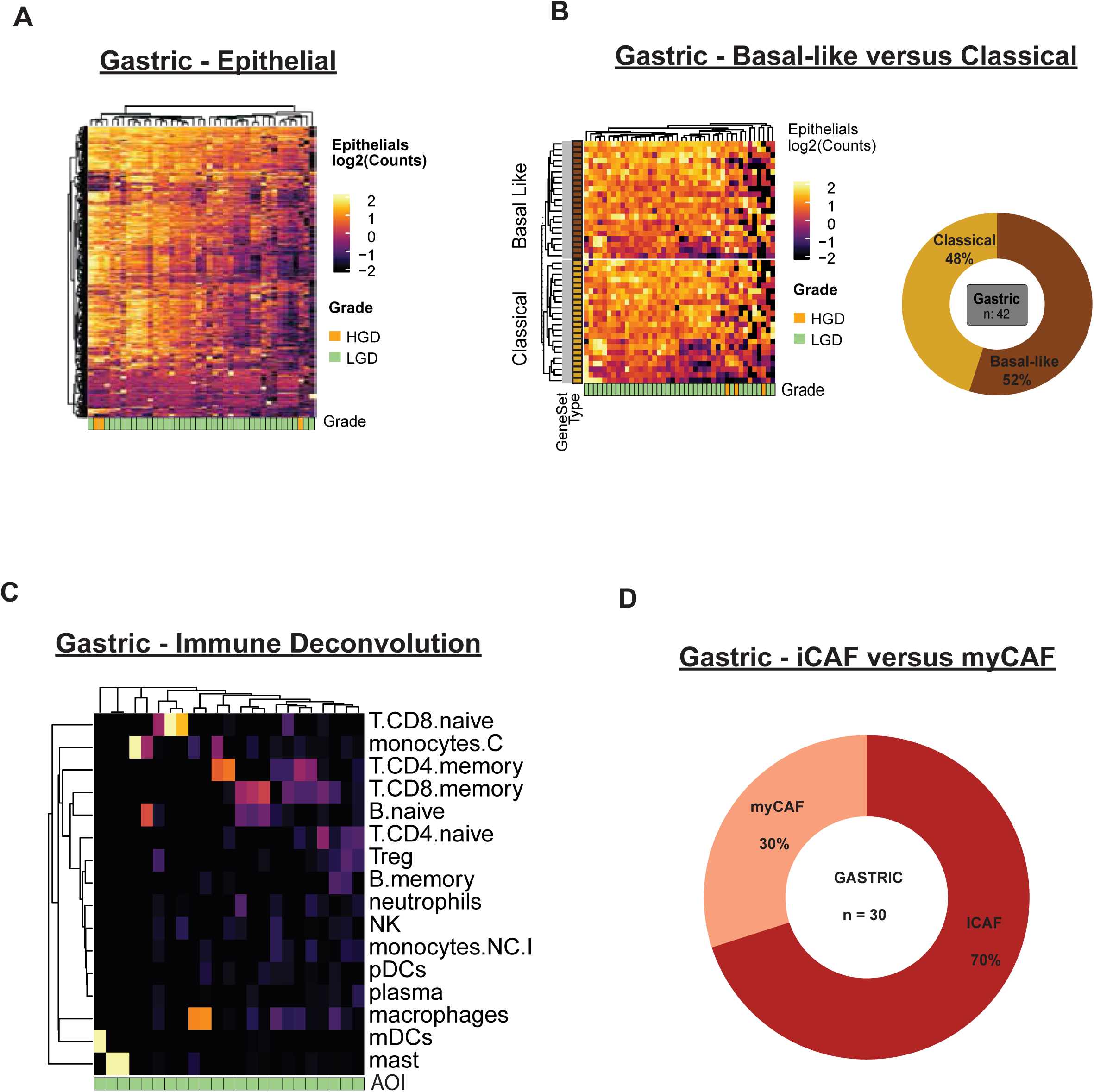
The Gastric subtype – a heterogenous, transitional cluster (A) Heatmap visualizing gene expression across all gastric epithelial Regions of Interest (ROIs) (n=42). The ROIs were subtyped into basal-like and classical epithelial phenotypes using established Moffitt gene signatures. Histological grade (HGD: LGD) is indicated below the heatmap columns. Unsupervised hierarchical clustering did not reveal distinct clusters based on histological grade. (B) Heatmap depicting the distribution of basal-like versus classical AOIs. Classification of gastric epithelial AOIs using the Moffitt metascore for each AOI identifies 49% as basal-like and 51% as Classical phenotypes. Pie chart showing the proportion of Gastric epithelial AOIs (n=42) classified as classical and basal-like. (C) Heatmap visualizing the relative enrichment of 16 distinct immune cell types across the gastric AOIs. The analysis revealed a diverse immune profile. (D) Donut chart showing the classification of gastric fibroblast AOIs (n=30) into subtypes using established metascores for inflammatory vs myofibroblastic cancer-associated fibroblast (CAF) populations (iCAF vs. myCAF). The Gastric subtype is predominantly enriched with the iCAF phenotype (70%) compared to the myCAF phenotype (30%).

### The Intestinal Subtype – A Classical Cluster

Unsupervised hierarchical clustering of all Intestinal epithelial AOIs did not reveal distinct clusters by grade (n=19) (**Figure 3A**). We used Moffitt gene signatures to calculate metascores in order to identify intestinal subtype-specific epithelial cell phenotypes. This analysis identifies that 95% of the intestinal AOIs are classical, and 5% are basal-like, revealing a predominantly classical phenotype on a heatmap (**Figure 3B**). To investigate the immune microenvironment of our intestinal cohort, we proceeded with an immune deconvolution analysis to determine the immune cell composition of the sub-type. Macrophages exhibited the highest relative enrichment, accounting for 28% of the total immune infiltrate, and CD4 and CD8 T memory cells representing 15% and 9% respectively (**Figure 3C**). Other immune cell populations, including B memory and naïve B cells showed lower representation. Meta score derived from myCAF and iCAF gene signatures annotated in PDAC to characterize the fibroblast niche. This analysis classified 60% of fibroblast AOIs in the intestinal subtype as iCAF and 40% as myCAF (**Figure 3D**). In summary, the intestinal subtype predominantly exhibits a classical epithelial cell phenotype and an inflammatory microenvironment enriched in macrophages and iCAF fibroblasts.

**Figure 3.**
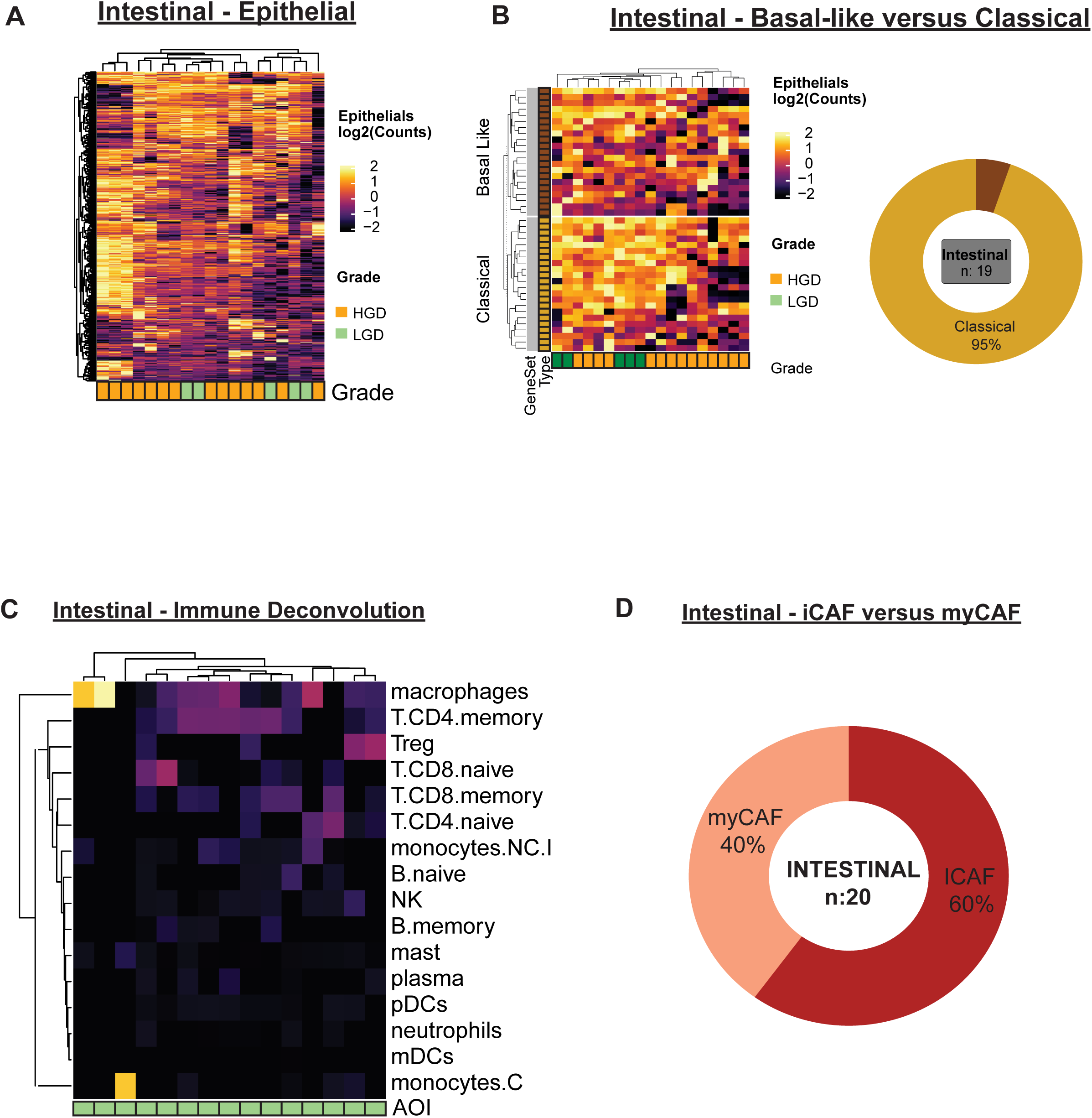
The Intestinal subtype – a classical, inflammatory Cluster (A) Heatmap visualizing gene expression across all intestinal epithelial Regions of Interest (ROIs) (n=19). The ROIs were subtyped into Basal-like and Classical epithelial phenotypes using established Moffitt gene signatures. Histological grade (HGD: LGD) is indicated below the heatmap columns. Unsupervised hierarchical clustering did not reveal distinct clusters based on histological grade. (B) Heatmap depicting the distribution of Basal-like versus Classical AOIs. Classification of intestinal epithelial AOIs using the Moffitt metascore for each AOI identifies 5% as basal-like and 95% as Classical phenotypes. Pie chart showing the proportion of intestinal epithelial AOIs (n=19) classified as classical and basal-like. (C) Heatmap visualizing the relative enrichment of 16 distinct immune cell types across the intestinal AOIs. The analysis revealed an enrichment of macrophages (D) Donut chart showing the classification of Intestinal fibroblast AOIs (n=20) into subtypes using established meta-scores for inflammatory vs myofibroblastic cancer-associated fibroblast (CAF) populations (iCAF vs. myCAF). The intestinal subtype is predominantly enriched with the iCAF phenotype (60%) compared to the myCAF phenotype (40%).

### The Pancreaticobiliary Subtype – A Basal-like, Immunosuppressed Cluster

Unsupervised hierarchical clustering of all PB epithelial AOIs (n=21) did not reveal distinct clusters by grade (**Figure 4A**). Again, we used Moffitt signatures to calculate metascores to identify PB subtype-specific epithelial cell phenotypes, this analysis classified 86% of epithelial AOIs in the PB subtype as Basal-like and 14% as Classical, revealing a predominantly basal-like signature on a heatmap (**Figure 4B**). To investigate the immune microenvironment of our PB cohort, we proceeded with an immune deconvolution analysis to determine the immune cell composition of the PB sub-type. Tregs exhibited the highest relative enrichment, accounting for 13% of the total immune infiltrate, with CD4 naïve T-cells and CD8 memory T-cells contributing 12% and 7% respectively (**Figure 4C**). Other immune cell populations, including macrophages and NK cells, showed lower representation. We used a metascore derived from previously defined myCAF and iCAF gene signatures annotated in PDAC [25, 26, 27] to characterize the fibroblast niche. This analysis classified 81% of fibroblast AOIs in the PB subtype as myCAF and 19% as iCAF (**Figure 4D**). In summary, the PB subtype predominantly exhibits a basal-like epithelial cell phenotype, and an immunosuppressive microenvironment enriched in Tregs and myCAF fibroblasts.

**Figure 4.**
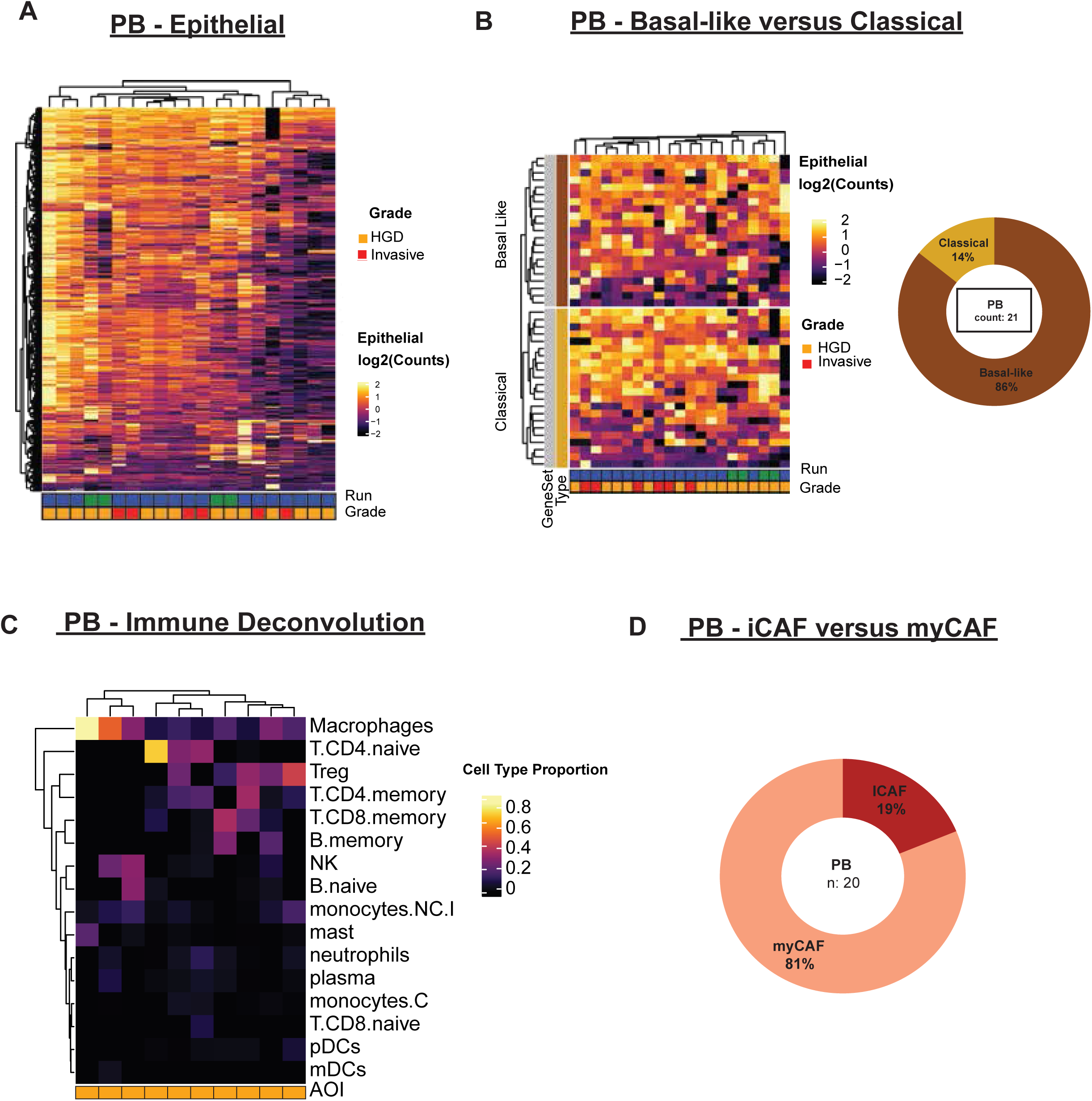
The Pancreaticobiliary Subtype - a Basal-Like, Immunosuppressed Cluster (A) Heatmap visualizing gene expression across all pancreatobiliary (PB) epithelial Regions of Interest (ROIs) (n=21). The ROIs were subtyped into basal-like and classical epithelial phenotypes using established Moffitt gene signatures. Histological grade (HGD: Invasive) is indicated below the heatmap columns. Unsupervised hierarchical clustering did not reveal distinct clusters based on histological grade. (B) Heatmap depicting the distribution of Basal-like versus Classical AOIs. Classification of PB epithelial AOIs using the Moffitt metascore for each AOI identifies 86% as basal-like and 14% as classical phenotypes. Pie chart showing the proportion of PB epithelial AOIs (n=21) classified as classical and basal-like. (C) Heatmap visualizing the relative enrichment of 16 distinct immune cell types across the PB AOIs. The analysis revealed an enrichment of T-regs. (D) Donut chart showing the classification of PB fibroblast AOIs (n=20) into subtypes using established meta-scores for inflammatory vs myofibrolastic cancer-Associated Fibroblast (CAF) populations (iCAF vs. myCAF). PB subtype is predominantly enriched with the iCAF phenotype (19%) compared to the myCAF phenotype (81%).

### The IPMN Transcriptomic Spectrum

To compare the transcriptomic range between IPMN subtypes, we identified the top 200 highly expressed genes in the gastric, intestinal and PB subtypes. Shared genes including the custom repeat genes were plotted against the median expression across the subtypes. The gastric subtype aligned closely with the median, the intestinal subtype showed overexpression, and the PB subtype exhibited an under-expression pattern (**Figure 5A**). Using the Moffitt meta score, we then observed a phenotypic spectrum with the Intestinal subtype being predominantly classical, the PB subtype being primarily basal-like, and the gastric subtype heterogeneous with near-equal representation of both phenotypes (**Figure 5B**). We then did a comparison of LGD and HGD, which did not reveal clear stratification in terms of classical versus basal-like phenotypes (**Figure 5C**). On comparing the Immune signatures between the subtypes, intestinal-type has a macrophage-enriched response, PB has a T-reg enriched response whereas gastric-type seems to have a diverse immune microenvironment (**Figure 5D**). Notably, analysis of *NKX6-2* in our cohort revealed significant enrichment in LGD IPMN compared to HGD (Median 65.4 vs 22.0; p = 4e-4) (**Figure 5E**). Using a fibroblast metascore, we then observed a phenotypic spectrum with the gastric subtype enriched in iCAFs, the PB subtype enriched in myCAF, and the Intestinal subtype heterogeneous with near-equal representation of both the CAF phenotypes (**Figure 5F**). To confirm the transitional and heterogeneous nature of gastric-type IPMN, we analyzed 11 additional FFPE whole resection specimens from 4 patients that were all identified as Gastric using the GeoMx Whole Transcriptomic Atlas panel. The Moffitt meta-score analysis of this cohort revealed significant intra-and inter-patient heterogeneity, with 5 out of 11 resections classified as basal-like and 6 out of 11 resections classified as classical, aligning with our earlier findings. (**Figure 5G**). These results suggest that IPMN represents a dynamic spectrum of cell phenotypes spanning between these histological subtypes.

**Figure 5.**
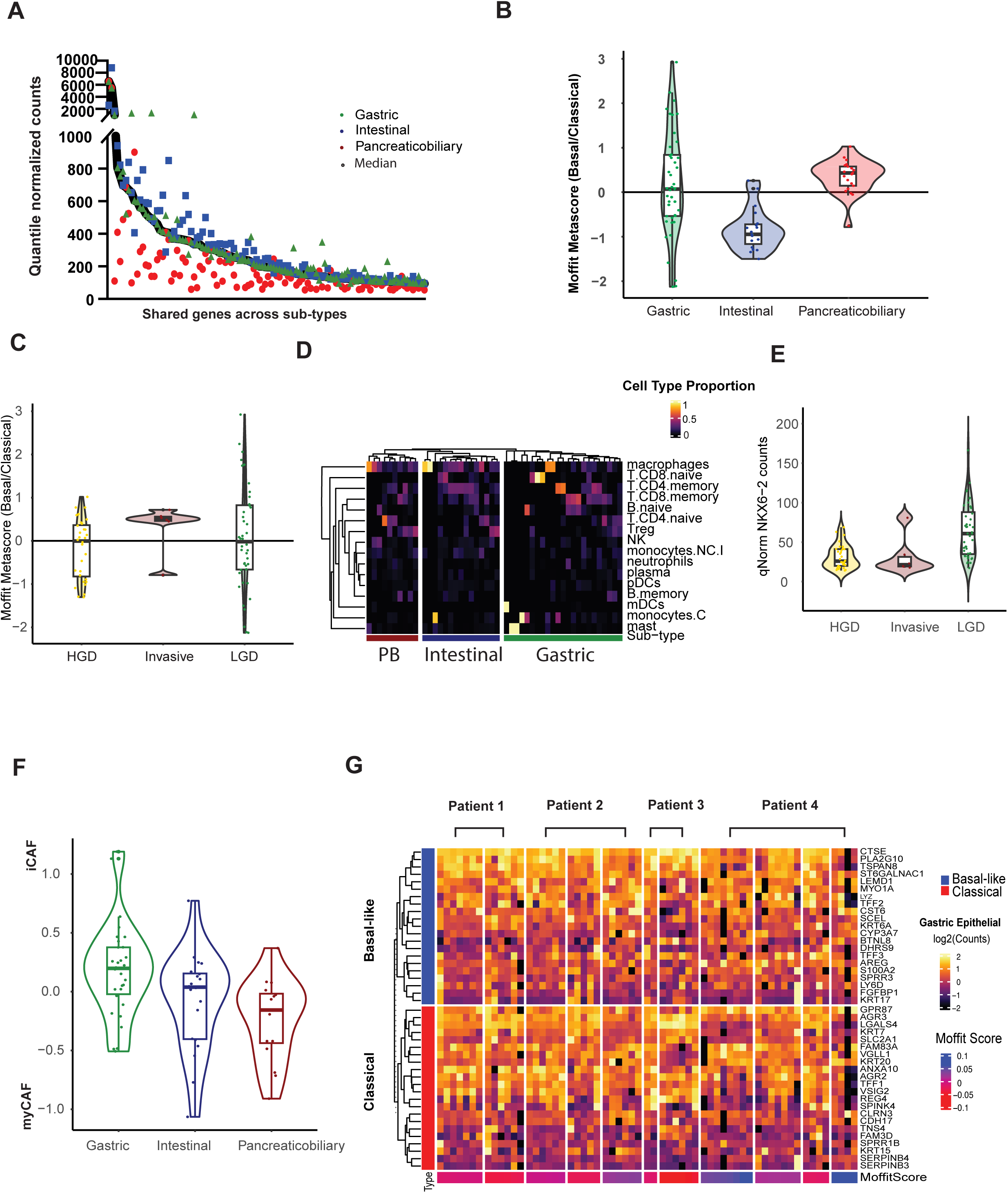
IPMN has a transcriptional spectrum across histological sub-types (A) Scatter plot of quantile-normalized counts for the top 200 highly expressed shared genes in the gastric, intestinal and pancreatobiliary (PB) subtypes, plotted against the median expression of these genes. The Gastric subtype aligns closely with the median expression, while the Intestinal subtype exhibits an overall over-expression pattern, and the PB subtype shows an under-expression pattern. (B) Violin plot of the Moffitt meta-score (basal-like/classical) comparing the three main IPMN subtypes. The intestinal subtype is predominantly classical-like, the PB subtype is primarily basal-like and the gastric subtype exhibits significant heterogeneity, centering close to zero. (C) Violin plot of the Moffitt meta-score comparing low-grade dysplasia (LGD), high-grade dysplasia (HGD), and invasive carcinoma. No clear stratification or significant shift between the basal-like and classical phenotypes is observed based on histological grade alone (D) Heatmap visualizing the relative proportions of 16 distinct immune cell types across the different epithelial subtypes. The intestinal subtype is characterized by a macrophage-enriched response, the PB subtype by a T-reg enriched response, and the gastric subtype displays a diverse and heterogeneous immune microenvironment. (E) Violin plot showing quantile-normalized NKX6-2gene expression counts. NKX6-2 expression is significantly higher in LGD compared to HGD and invasive carcinoma. (F) Violin plot of the cancer-associated fibroblast (CAF) meta-score (myCAF/iCAF) across IPMN subtypes. The gastric subtype is primarily enriched in the inflammatory CAF (iCAF) phenotype, the PB subtype is enriched in the myofibroblastic CAF (myCAF) phenotype, and the intestinal subtype shows a heterogeneous representation of both phenotypes. (G) Heatmap of key epithelial subtyping genes and the corresponding Moffitt meta-score classification in 11 additional FFPE whole resection specimens from 4 independent patients (P1-P4). The analysis confirms the heterogeneous nature of gastric IPMN, with resections classifying as both basal-like (5/11) and classical (6/11) states.

### Repeat Element Dysregulation in the IPMN Spectrum

To investigate the heterogeneous landscape of IPMN, we analyzed the RNA expression of retrotransposons, including long interspersed nuclear element 1 (LINE1) retrotransposon open reading frame 1 (ORF1) and open reading frame 2 (ORF2), HSATII, and two human endogenous retroviruses (HERV-K and HERV-H), across the different IPMN subtypes.

Repetitive elements emerged as some of the most highly expressed genes in the epithelial compartment of gastric, intestinal and PB subtypes and we found that their expression levels were significantly higher than KRT7 in IPMN (**Figure 6A**). Between the different subtypes, median LINE1 ORF1 was significantly higher in intestinal (median - 500.67; padj = 0.002) and PB (median - 458.37; p=0.02) compared to gastric subtype (median - 346.36). LINE1 ORF2 was also significantly higher in intestinal (median = 869.68; padj = 0.002) and PB (median = 784.48; p=0.02) compared to gastric subtype (median = 601.14). HSAT II was significantly higher in intestinal (median = 6860.77; padj = 0.01) and PB (median = 7251.57; p=0.001) compared to gastric (median = 4515.87). HERVK was significantly higher in Intestinal (median = 17.62; padj = 0.01) compared to gastric subtype (median = 14.68). HERVH was significantly higher in intestinal (median = 16.83; padj = 0.04) compared to gastric (median = 14.46) (**Figure 6B**). Repeat expression was higher in IPMN with HGD compared to LGD for HSATII (Median 7440 vs 4487; p = 0.0001), LINE1 ORF1 (Median 487 vs 351; p = 0.001), and LINE1 ORF2 (Median 839 vs 607; p = 0.001) (**Figure 6C**). These findings indicate that repetitive element dysregulation is enriched in HGD with associated intestinal and PB histology.

**Figure 6:**
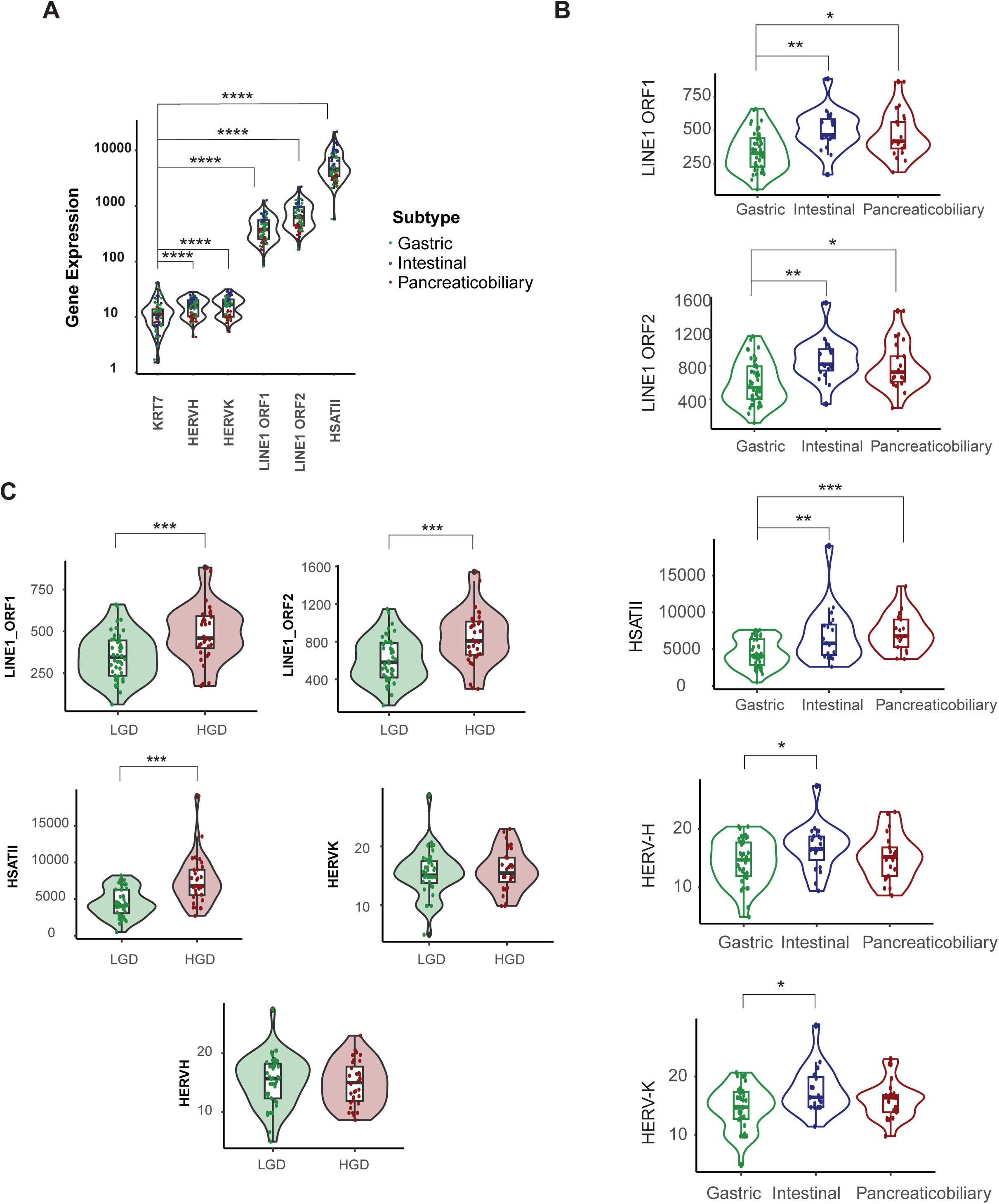
Repeat Element Expression in IPMN Phenotypes (A) Violin plots comparing the quantile-normalized gene expression of the epithelial marker KRT7 against key repetitive elements: LINE1 ORF1, LINE1 ORF2, HSATII, HERV-H, HERV-K across all IPMN epithelial regions. The repetitive elements are among the most highly expressed genes in the cohort, with significantly higher expression levels than the epithelial marker KRT7. (B) Violin plots comparing the expression of five repetitive elements across the gastric (green), intestinal (blue), and pancreaticobiliary (PB, red) subtypes. (C) Violin plots comparing the expression of five repetitive elements across in low-grade dysplasia (LGD) vs high grade dysplasia (HGD.

### Single-cell Transcriptomic Analysis of the IPMN Spectrum

To comprehensively characterize cell phenotype variations between different histological subtypes of IPMN at the single-cell level, we employed the NanoString CosMx Spatial Molecular Imager (SMI) platform. This approach utilized a 1,000-plex RNA panel with custom probes targeting long interspersed nuclear element-1 (LINE1) retrotransposon open reading frames (ORF1 and ORF2), HSATII, and two human endogenous retroviruses (HERV-K and HERV-H), enabling the in-situ visualization of individual RNA molecules. Spatially resolved single-cell transcriptomic data were collected from 50 IPMN resections across 17 patients, encompassing 217,821 cells imaged from 359 fields of view (FOVs) (**Figure 7A, 7B**). Uniform Manifold Approximation and Projection (UMAP) of epithelial cell molecular signatures demonstrated clear separation based on both pathological grade and histological subtype where cells segregated progressively along the malignancy spectrum, with HGD, HGD near an invasive focus and invasive carcinoma, forming a distinct cluster away from LGD and normal cells (**Figure 7C**). The intestinal and PB subtypes had two distinct clusters, but there was notable mixing of all three histological subtypes in the largest clusters **(Figure 7D).** Pseudotime trajectory analysis was employed to model the temporal progression of cellular states within each subtype **(Figure 7E-G).** In all three subtypes, the trajectory initiates from primarily gastric state with overlapping intestinal and PB cells **(Figure 7E and F)**. This primary starting cluster then branches out to distinct intestinal and PB clusters with increased HGD and invasive disease (**Figure 7G**). This suggests that malignant progression follows a subtype-specific transcriptional pathway starting from a common gastric enriched state. Furthermore, mapping classical and basal states using the Moffitt signature we find increased classical phenotype in intestinal clusters, but interestingly a basal and classical cluster in the PB subtype (**Figure 7H).** Repeat RNA expression was also observed to be higher in advanced HGD lesions in the intestinal and PB subtype, compared to gastric-type IPMN (**Figure 7I**). Collectively, the single cell spatial analysis reveals that IPMN subtypes start from a gastric cell state with distinct trajectories towards intestinal and PB subtypes with increasing dysplasia and aberrant repeat RNA expression.

**Figure 7.**
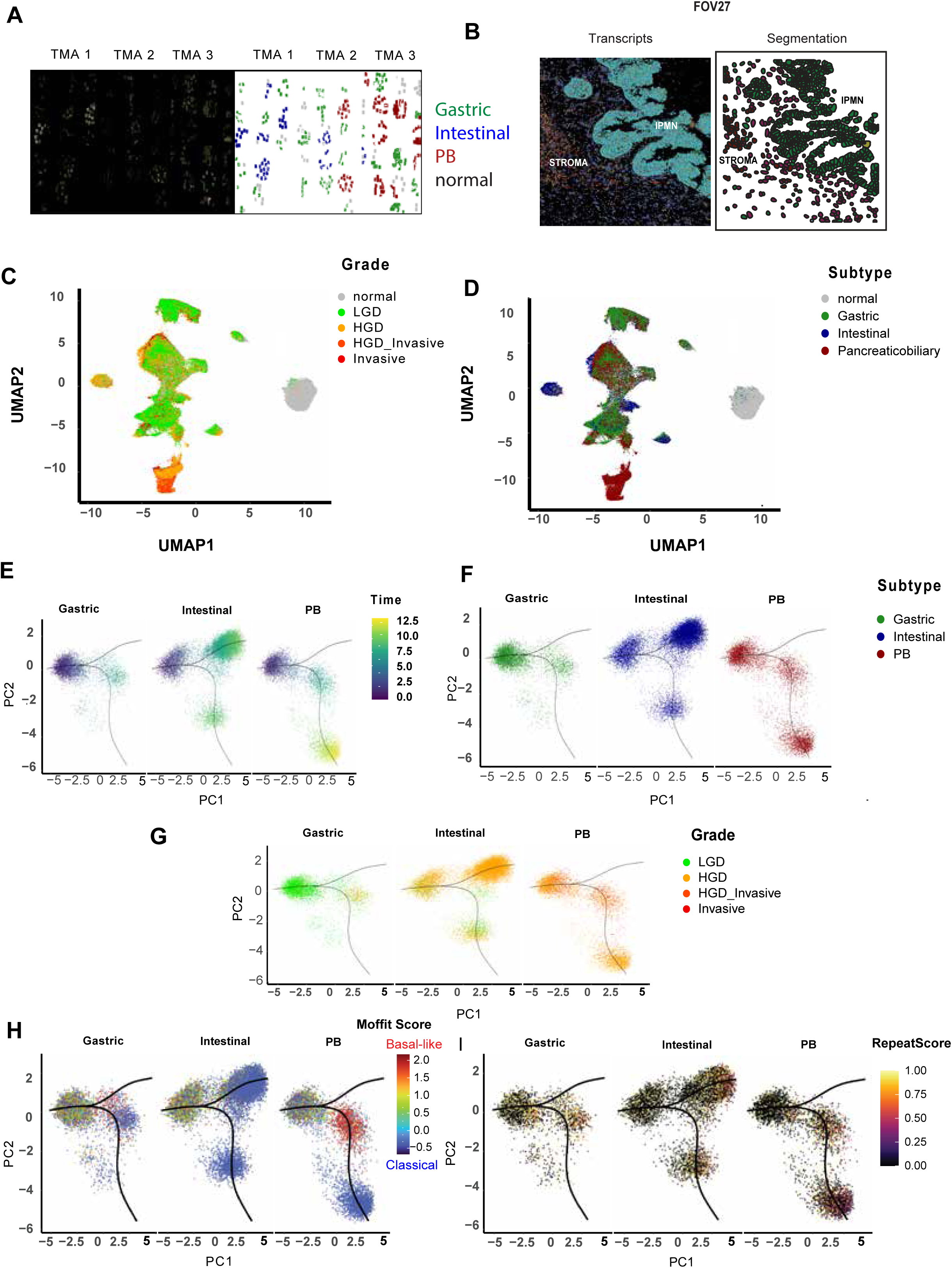
Single-Cell Spatial Transcriptomic Analysis of IPMN Subtypes (A) Fluorescence images (left) and corresponding single-cell transcript segmentation and transcript visualization (right) for epithelial cells across three representative TMA cores using the CosMx Spatial Molecular Imager (SMI) platform. Data were collected from 50 IPMN resections across 17 patients, encompassing 217,821 cells from 359 fields of view (FOV). (B) Representative FOVs illustrating the in-situ visualization of individual RNA molecules (left) and the subsequent segmentation of epithelial (IPMN) cells from the surrounding stroma (right). A 1,000-plex RNA panel, including custom probes for repetitive elements was used. (C) Uniform Manifold Approximation and Projection (UMAP) of epithelial cell molecular signatures colored by pathological grade. Cells segregate progressively along the malignancy spectrum, with high-grade dysplasia (HGD) and invasive lesions near an invasive focus forming a distinct cluster, demonstrating molecular separation from low-grade dysplasia (LGD) and normal cells. (D) UMAP analysis colored by histological subtype gastric, intestinal and pancreaticobiliary (PB). While Intestinal and PB cells show distinct clustering, there is significant overlap and mixing of all three subtypes along with gastric in the largest molecular clusters. (E) Principal Component (PC) plots showing the Pseudotime trajectory of all epithelial cells, separated by subtype. Trajectories for all three subtypes initiate from a common state and show increasing Pseudotime (“Time”) along the divergence vector. (F) Principal Component (PC) plots showing the Pseudotime trajectory of all epithelial cells, separated by subtype. The initial starting cluster is predominantly composed of Gastric cells diverging into 2 trajectories enriched by intestinal and PB subtypes. (G) The trajectory branches to distinct Intestinal and PB clusters, correlating with increasing HGD and Invasive disease, suggesting a subtype-specific transcriptional progression from a common gastric-enriched cell state. (H) PC plots showing the distribution of the basal-like (red) and classical (blue) phenotypes (Moffitt Score) onto the subtype trajectories. The intestinal trajectory is largely classical, while the PB subtype shows a distinct basal-like enrichment. (I) PC plots showing the normalized Repeat Score onto the subtype trajectories. High Repeat Score is concentrated in the advanced HGD and Invasive lesions of the intestinal and PB subtype trajectories.

## DISCUSSION

The clinical dilemma of managing patients with IPMN has led to a variety of studies focused on identifying biomarkers of IPMN progression to invasive carcinoma. This current work using spatial transcriptomic technologies has validated findings from other studies and contributes new analyses involving repeat RNA characterization linked with changes in morphology and dysplasia. The combination of focal whole transcriptome and single cell trajectory analysis has provided support of a molecular continuum of IPMN starting from low-grade gastric morphology with progression to a PB or intestinal subtype with increasing dysplasia.

The current work has been consistent with other studies of the IPMN immune microenvironment with a diverse immune microenvironment in gastric IPMN including plasma cells as seen by Sans et al. [31] and infiltration of macrophages with HGD consistent with Eckhoff et al. [30]. Notably, our analysis of CAFs reveals a shift from iCAF to myCAF phenotype with the progression from a gastric to higher grade intestinal and PB subtypes **(Figure 5).** To obtain a more comprehensive view of the cellular ecosystem of IPMN, recent studies have utilized spatial transcriptomic technologies [30, 31, 32]. Eckhoff et al. [30] used the Bruker/Nanostring GeoMx cancer transcriptome atlas of approximately 1800 genes on 11 IPMN specimens and found that the immune infiltrate of IPMN with LGD are enriched for T-cells, while IPMN with HGD had a predominance of macrophage infiltrates. Although epithelial and fibroblast areas of interest (AOIs) were obtained, they were not the focus of this initial spatial transcriptomic study of IPMN. Sans et al. [31] used the Visium high density spot assay of 8000 genes on 13 snap-frozen IPMN samples. This study had 12,685 AOIs comprised of mixed cell populations and utilized a robust cell-type deconvolution method to digitally separate cell populations. Separating samples based on the degree of dysplasia, the authors identified gastric isthmus-pit (GIP) phenotype with high *NKX6-2* in IPMN with LGD and associated plasma cell infiltrates, while elevated *CLDN4* and *CEACAM6* was seen in IPMN with HGD and invasive carcinoma. Loss of NKX6-2 was also associated with increased epithelial-mesenchymal transition (EMT) markers consistent with cell state changes toward invasive cancer. Agostini et al. [32] utilized Visium in 18 IPMN samples from 14 patients and GeoMx whole transcriptome atlas (18,676 genes) in 57 IPMN samples from 40 patients. This work also found higher *NKX6-2* correlating with gastric isthmus markers and identified *HOXB3* and *ZNF117* as markers for LGD. We evaluated the previously reported IPMN dysplasia gene *NKX6-2* in our cohort and identified significant enrichment in LGD compared to HGD. This is consistent with Sans et al. [31], and supports *NKX6-2* as an important marker of IPMN gastric LGD state that merits further study.

Repeat RNA expression in the context of PDAC cells was shown to be associated with EMT [33], which we found in IPMN to be more increased in higher-grade intestinal and PB cell-states, compared to the lower grade gastric lesions (**Figure 6**). This indicates that dysplasia and associated repeat dysregulation can occur in either epithelial or mesenchymal/basal-like states. This relationship of morphology, EMT cell state, and repeat expression became more evident with single-cell spatial analysis revealing dysregulated repeats associated cell state changes to intestinal and PB phenotype with either classical epithelial or basal-like transcriptional programs (**Figure 7**). Altogether, these data support repeat RNA expression as a marker of cellular transdifferentiation and dysplasia that can serve as a risk stratification biomarker independent of morphology. These repeat RNAs can be used to complement other IPMN risk stratification markers that have been identified, including the Das-1 antibody [34, 35, 36], mutation-based assessment [37], and other multimodality approaches [38, 39, 40]. In summary, the collective spatial transcriptomics studies have refined our understanding of IPMN progression to PDAC that has opened a new era of precision oncology to identify biomarkers for risk stratification and implications for therapeutic cancer interception.

Despite these advancements, our study has several limitations, including the retrospective nature and selection bias inherent to a cohort consisting solely of resected IPMN lesions.

Furthermore, while the pseudotime analysis strongly supports a molecular continuum, it represents a transcriptional distance rather than direct proof of in vivo temporal progression. Future efforts will focus on validating the translational utility of the identified markers, such as repeat RNAs in clinical specimens.

## FUNDING

Dana Farber/Harvard Cancer Center: GI SPORE 5P50CA127003-12 Career Development Award (YH-B)

MGH Executive Committee on Research Center for Diversity and Inclusion Physician Scientist Award (YH-B)

National Institutes of Health grant U01CA228963 (DTT), NIH/NCI Diversity Supplement: R01 CA215498-02 (YH-B)

Massachusetts Life Sciences Center Research Infrastructure Program (DTT)

## ACKNOWLEDGEMENTS

We are grateful to Danielle Bestoso and Angelique N. Gilbert for administrative support. Funding support from:

## AUTHOR CONTRIBUTIONS

Conceptualization AP, MJR, MLZ, DTT, MM-K, YGH-B Formal Analysis AP, MJR, MLZ, DTT, YGH-B

Investigation AP, MJR, MLZ, YS, BKP, KX, JRK, NJ-C, SM, EL,LTN, MLZ, NB, VD, MJA, MM-K, DTT, YGH-B

Methodology AP, MJR, MLZ, YS, BKP, KX, NP, LTN, MM-K, VD, MJA, DTT, YGH-B Resources LTN, NB, VD, CF-D, MM-K

Writing AP, MJR, MLZ, MM-K, DTT, YGH-B

Visualization AP, MJR, YS, BKP, KX, MJA, DTT Supervision AP, MLZ, LTN, MJA, MM-K, DTT, YGH-B

Project Administration AP, MJA, MM-K, DTT, YGH-B Funding Acquisition DTT, YGH-B

## DATA AVAILABILITY STATEMENT

All spatial transcriptomic data will be uploaded to NCBI GEO or other appropriate public repository. Otherwise, all data will be provided upon request. All analysis codes will be provided upon request.

## REFERENCES

1 Farrell JJ, Fernandez-del Castillo C. Pancreatic cystic neoplasms: management and unanswered questions. Gastroenterology 2013;144:1303–15.

2 Rech AJ, Vonderheide RH. Dynamic interplay of oncogenes and T cells induces PD-L1 in the tumor microenvironment. Cancer Discov 2013;3:1330–2.

3 Ohtsuka T, Fernandez-Del Castillo C, Furukawa T, Hijioka S, Jang JY, Lennon AM, et al. International evidence-based Kyoto guidelines for the management of intraductal papillary mucinous neoplasm of the pancreas. Pancreatology 2024;24:255–70.

4 Roldan J, Harrison JM, Qadan M, Bolm L, Baba T, Brugge WR, et al. “Evolving Trends in Pancreatic Cystic Tumors: A 3-Decade Single-Center Experience With 1290 Resections”. Ann Surg 2023;277:491–7.

5 Gardner TB, Park WG, Allen PJ. Diagnosis and Management of Pancreatic Cysts. Gastroenterology 2024;167:454–68.

6 Wood LD, Canto MI, Jaffee EM, Simeone DM. Pancreatic Cancer: Pathogenesis, Screening, Diagnosis, and Treatment. Gastroenterology 2022;163:386–402.e1.

7 Bayne LJ, Beatty GL, Jhala N, Clark CE, Rhim AD, Stanger BZ, et al. Tumor-derived granulocyte-macrophage colony-stimulating factor regulates myeloid inflammation and T cell immunity in pancreatic cancer. Cancer Cell 2012;21:822–35.

8 Beatty GL, Winograd R, Evans RA, Long KB, Luque SL, Lee JW, et al. Exclusion of T Cells From Pancreatic Carcinomas in Mice Is Regulated by Ly6C(low) F4/80(+) Extratumoral Macrophages. Gastroenterology 2015;149:201–10.

9 Chang JH, Jiang Y, Pillarisetty VG. Role of immune cells in pancreatic cancer from bench to clinical application: An updated review. Medicine (Baltimore) 2016;95:e5541.

10 Clark CE, Hingorani SR, Mick R, Combs C, Tuveson DA, Vonderheide RH. Dynamics of the immune reaction to pancreatic cancer from inception to invasion. Cancer Res 2007;67:9518–27.

11 Erkan M, Hausmann S, Michalski CW, Fingerle AA, Dobritz M, Kleeff J, et al. The role of stroma in pancreatic cancer: diagnostic and therapeutic implications. Nat Rev Gastroenterol Hepatol 2012;9:454–67.

12 Jang JE, Hajdu CH, Liot C, Miller G, Dustin ML, Bar-Sagi D. Crosstalk between Regulatory T Cells and Tumor-Associated Dendritic Cells Negates Anti-tumor Immunity in Pancreatic Cancer. Cell Rep 2017;20:558–71.

13 Marks DL, Olson RL, Fernandez-Zapico ME. Epigenetic control of the tumor microenvironment. Epigenomics 2016;8:1671–87.

14 Moffitt RA, Marayati R, Flate EL, Volmar KE, Loeza SG, Hoadley KA, et al. Virtual microdissection identifies distinct tumor-and stroma-specific subtypes of pancreatic ductal adenocarcinoma. Nat Genet 2015;47:1168–78.

15 Patra KC, Kato Y, Mizukami Y, Widholz S, Boukhali M, Revenco I, et al. Mutant GNAS drives pancreatic tumourigenesis by inducing PKA-mediated SIK suppression and reprogramming lipid metabolism. Nat Cell Biol 2018;20:811–22.

16 Rooney MS, Shukla SA, Wu CJ, Getz G, Hacohen N. Molecular and genetic properties of tumors associated with local immune cytolytic activity. Cell 2015;160:48–61.

17 Timosenko E, Hadjinicolaou AV, Cerundolo V. Modulation of cancer-specific immune responses by amino acid degrading enzymes. Immunotherapy 2017;9:83–97.

18 Zhang Y, Yan W, Mathew E, Bednar F, Wan S, Collins MA, et al. CD4+ T lymphocyte ablation prevents pancreatic carcinogenesis in mice. Cancer Immunol Res 2014;2:423–35.

19 Zhang Y, Zoltan M, Riquelme E, Xu H, Sahin I, Castro-Pando S, et al. Immune Cell Production of Interleukin 17 Induces Stem Cell Features of Pancreatic Intraepithelial Neoplasia Cells. Gastroenterology 2018;155:210–23 e3.

20 Zhu Y, Herndon JM, Sojka DK, Kim KW, Knolhoff BL, Zuo C, et al. Tissue-Resident Macrophages in Pancreatic Ductal Adenocarcinoma Originate from Embryonic Hematopoiesis and Promote Tumor Progression. Immunity 2017;47:597.

21 Roth S, Zamzow K, Gaida MM, Heikenwälder M, Tjaden C, Hinz U, et al. Evolution of the immune landscape during progression of pancreatic intraductal papillary mucinous neoplasms to invasive cancer. EBioMedicine 2020;54:102714.

22 Hernandez S, Parra ER, Uraoka N, Tang X, Shen Y, Qiao W, et al. Diminished Immune Surveillance during Histologic Progression of Intraductal Papillary Mucinous Neoplasms Offers a Therapeutic Opportunity for Cancer Interception. Clin Cancer Res 2022;28:1938–47.

23 Mo S, Zou L, Hu Y, Chang X, Chen J. Expression of PD-L1 and VISTA in Intraductal Papillary Mucinous Neoplasm With Associated Invasive Carcinoma of the Pancreas. Mod Pathol 2023;36:100223.

24 Bernard V, Semaan A, Huang J, San Lucas FA, Mulu FC, Stephens BM, et al. Single-Cell Transcriptomics of Pancreatic Cancer Precursors Demonstrates Epithelial and Microenvironmental Heterogeneity as an Early Event in Neoplastic Progression. Clin Cancer Res 2019;25:2194–205.

25 Biffi G, Oni TE, Spielman B, Hao Y, Elyada E, Park Y, et al. IL1-Induced JAK/STAT Signaling Is Antagonized by TGFbeta to Shape CAF Heterogeneity in Pancreatic Ductal Adenocarcinoma. Cancer Discov 2019;9:282–301.

26 Elyada E, Bolisetty M, Laise P, Flynn WF, Courtois ET, Burkhart RA, et al. Cross-Species Single-Cell Analysis of Pancreatic Ductal Adenocarcinoma Reveals Antigen-Presenting Cancer-Associated Fibroblasts. Cancer Discov 2019;9:1102–23.

27 Ohlund D, Elyada E, Tuveson D. Fibroblast heterogeneity in the cancer wound. J Exp Med 2014;211:1503–23.

28 Kakizaki Y, Makino N, Tozawa T, Honda T, Matsuda A, Ikeda Y, et al. Stromal Fibrosis and Expression of Matricellular Proteins Correlate With Histological Grade of Intraductal Papillary Mucinous Neoplasm of the Pancreas. Pancreas 2016;45:1145–52.

29 Shindo K, Aishima S, Ohuchida K, Fujino M, Mizuuchi Y, Hattori M, et al. Podoplanin expression in the cyst wall correlates with the progression of intraductal papillary mucinous neoplasm. Virchows Arch 2014;465:265–73.

30 Eckhoff AM, Fletcher AA, Landa K, Iyer M, Nussbaum DP, Shi C, et al. Multidimensional Immunophenotyping of Intraductal Papillary Mucinous Neoplasms Reveals Novel T Cell and Macrophage Signature. Ann Surg Oncol 2022;29:7781–8.

31 Sans M, Makino Y, Min J, Rajapakshe KI, Yip-Schneider M, Schmidt CM, et al. Spatial Transcriptomics of Intraductal Papillary Mucinous Neoplasms of the Pancreas Identifies NKX6-2 as a Driver of Gastric Differentiation and Indolent Biological Potential. Cancer Discov 2023;13:1844–61.

32 Agostini A, Piro G, Inzani F, Quero G, Esposito A, Caggiano A, et al. Identification of spatially-resolved markers of malignant transformation in Intraductal Papillary Mucinous Neoplasms. Nat Commun 2024;15:2764.

33 You E, Danaher P, Lu C, Sun S, Zou L, Phillips IE, et al. Disruption of cellular plasticity by repeat RNAs in human pancreatic cancer. Cell 2024.

34 Das KK, Brown JW, Fernandez Del-Castillo C, Huynh T, Mills JC, Matsuda Y, et al. mAb Das-1 identifies pancreatic ductal adenocarcinoma and high-grade pancreatic intraepithelial neoplasia with high accuracy. Hum Pathol 2021;111:36–44.

35 Das KK, Geng X, Brown JW, Morales-Oyarvide V, Huynh T, Pergolini I, et al. Cross Validation of the Monoclonal Antibody Das-1 in Identification of High-Risk Mucinous Pancreatic Cystic Lesions. Gastroenterology 2019;157:720–30 e2.

36 Das KK, Xiao H, Geng X, Fernandez-Del-Castillo C, Morales-Oyarvide V, Daglilar E, et al. mAb Das-1 is specific for high-risk and malignant intraductal papillary mucinous neoplasm (IPMN). Gut 2014;63:1626–34.

37 Springer S, Wang Y, Dal Molin M, Masica DL, Jiao Y, Kinde I, et al. A combination of molecular markers and clinical features improve the classification of pancreatic cysts. Gastroenterology 2015;149:1501–10.

38 Brugge WR, Lewandrowski K, Lee-Lewandrowski E, Centeno BA, Szydlo T, Regan S, et al. Diagnosis of pancreatic cystic neoplasms: a report of the cooperative pancreatic cyst study. Gastroenterology 2004;126:1330–6.

39 Springer S, Masica DL, Dal Molin M, Douville C, Thoburn CJ, Afsari B, et al. A multimodality test to guide the management of patients with a pancreatic cyst. Sci Transl Med 2019;11.

40. Kwan, M. C., Pitman, M. B., Fernandez-Del Castillo, C., & Zhang, M. L. (2024). Revisiting the performance of cyst fluid carcinoembryonic antigen as a diagnostic marker for pancreatic mucinous cysts: a comprehensive 20-year institutional review. Gut, 73(4), 629–638.

