## Supplementary Table 1 for "Intraductal Papillary Mucinous Neoplasm Cellular Plasticity is Linked with Repeat Element Dysregulation"

| **Supplementary Table 1 : Patient cohort and Demographics : IPMN Tissue microarray (TMA)** | | | | | |  |
| --- | --- | --- | --- | --- | --- | --- |
| **Case ID** | **Slide number** | **TMA number** | **Age (decade)** | **Sex** | **Subtype** | **Slide grade** |
| Case 1 | 1A | TMA 2 | 80s | M | gastric | LGD |
|  | 1B | TMA 2 | 80s | M | gastric | LGD |
| Case 2 | 2A | TMA 3 | 70s | M | PB | HGD |
|  | 2B | TMA 3 | 70s | M | gastric | LGD |
|  | 2C | TMA 3 | 70s | M | intestinal | HGD |
| Case 3 | 3A | TMA 2 | 70s | M | gastric | LGD |
| Case 4 | 4A | TMA 2 | 70S | F | gastric | LGD |
|  | 4B | TMA 2 | 70S | F | gastric | HGD |
|  | 4C | TMA 2 | 70S | F | gastric | LGD |
| Case 5 | 5A | TMA 1 | 50s | F | intestinal | HGD |
|  | 5B | TMA 1 | 50s | F | intestinal | HGD |
| Case 6 | 6A | TMA 1 | 50s | F | intestinal | HGD |
|  | 6B | TMA 1 | 50s | F | intestinal | HGD |
| Case 7 | 7A | TMA 1 | 60s | M | gastric | LGD |
|  | 7B | TMA 1 | 60s | M | gastric | LGD |
| Case 8 | 8A | TMA 1 | 60s | M | gastric | LGD |
|  | 8B | TMA 1 | 60s | M | gastric | HGD |
|  | 8C | TMA 1 | 60s | M | gastric | LGD |
| Case 9 | 9A | TMA 2 | 60s | F | intestinal | HGD |
|  | 9B | TMA 2 | 60s | F | intestinal | LGD |
|  | 9C | TMA 2 | 60s | F | intestinal | HGD |
|  | 9D | TMA 2 | 60s | F | intestinal | HGD |
|  | 9E | TMA 2 | 60s | F | intestinal | LGD |
|  | 9F | TMA 2 | 60s | F | gastric | LGD |
| Case 10 | 10A | TMA 1 | 60s | F | intestinal | HGD |
|  | 10B | TMA 1 | 60s | F | gastric | LGD |
|  | 10C | TMA 1 | 60s | F | intestinal | LGD |
|  | 10D | TMA 1 | 60s | F | intestinal | HGD |
|  | 10E | TMA 1 | 60s | F | intestinal | LGD |
| Case 11 | 11A | TMA 1 | 70s | M | gastric | LGD |
|  | 11B | TMA 1 | 70s | M | gastric | HGD |
| Case 12 | 12A | TMA 3 | 60s | M | PB | HGD |
|  | 12B | TMA 3 | 60s | M | PB | HGD |
|  | 12C | TMA 3 | 60s | M | PB | HGD |
|  | 12D | TMA 3 | 60s | M | PB | Invasive |
|  | 12E | TMA 3 | 60s | M | PB | HGD |
|  | 12F | TMA 3 | 60s | M | PB | Invasive |
| Case 13 | 13A | TMA 2 | 70s | M | gastric | LGD |
|  | 13B | TMA 2 | 70s | M | PB | HGD |
|  | 13C | TMA 2 | 70s | M | gastric | LGD |
| Case 14 | 14A | TMA 3 | 70s | M | PB | HGD |
|  | 14B | TMA 3 | 70s | M | gastric | LGD |
|  | 14C | TMA 3 | 70s | M | gastric | LGD |
|  | 14D | TMA 3 | 70s | M | PB | invasive |
|  | 14E | TMA 3 | 70s | M | gastric | LGD |
| Case 15 | 15A | TMA 1 | 80s | F | gastric | LGD |
|  | 15B | TMA 1 | 80s | F | gastric | LGD |
| Case 16 | 16A | TMA 3 | 50s | F | gastric | LGD |
|  | 16B | TMA 3 | 50s | F | gastric | LGD |
| Case 17 | 17A | TMA 2 | 70s | F | PB | HGD |
|  | 17B | TMA 2 | 70s | F | gastric | LGD |
| Case 18 (not captured on GeoMx) | 18A | TMA 3 | 70s | M | PB | HGD |
