## Supplementary Table 2 for "Intraductal Papillary Mucinous Neoplasm Cellular Plasticity is Linked with Repeat Element Dysregulation"

| **Supplementary Table 2 : Patient cohort and Demographics: IPMN whole slide resections** | | | | | |
| --- | --- | --- | --- | --- | --- |
| **Case number** | **Slide number** | **Age (decade)** | **Sex** | **Subtype** | **Slide grade** |
| Case 1 | 1A | 60s | F | Gastric | LGD |
|  | 1B | 60s | F | Gastric | LGD |
| Case 2 | 2A | 60s | F | Gastric | LGD |
|  | 2B | 60s | F | Gastric | LGD |
| Case 3 | 3A | 70s | M | Gastric | LGD |
|  | 3B | 70s | M | Gastric | LGD |
|  | 3C | 70s | M | Gastric | HGD |
| Case 4 | 4A | 80s | F | Gastric | HGD |
|  | 4B | 80s | F | Gastric | HGD |
|  | 4C | 80s | F | Gastric | LGD |
|  | 4D | 80s | F | Gastric | LGD |
